# Experimental and statistical re-evaluation provides no evidence for *Drosophila* courtship song rhythms

**DOI:** 10.1101/140483

**Authors:** David L. Stern, Jan Clemens, Philip Coen, Adam J. Calhoun, John B. Hogenesch, Ben Arthur, Mala Murthy

## Abstract

From 1980 to 1992, a series of influential papers reported on the discovery, genetics, and evolution of a periodic cycling of the interval between *Drosophila* male courtship song pulses. The molecular mechanisms underlying this periodicity were never described. To reinitiate investigation of this phenomenon, we performed automated segmentation of songs, but failed to detect the proposed periodicity [Arthur BJ et al. (2013) *BMC Biol* 11:11; Stern DL (2014) *BMC Biol* 12:38]. Kyriacou CP et al. [(2017) *PNAS* 114:1970-1975] report that we failed to detect song rhythms because i) our flies did not sing enough and ii) our segmenter did not identify many of the song pulses. Kyriacou et al. manually annotated a subset of our recordings and reported that two strains displayed rhythms with genotype-specific periodicity, in agreement with their original reports. We cannot replicate this finding and show that the manually-annotated data, the original automatically segmented data, and a new data set provide no evidence for either the existence of song rhythms or song periodicity differences between genotypes. Furthermore, we have re-examined our methods and analysis and find that our automated segmentation method was not biased to prevent detection of putative song periodicity. We conclude that there is currently no evidence for the existence of *Drosophila* courtship song rhythms.

**Significance statement:** Previous studies have reported that male vinegar flies sing courtship songs with a periodic rhythm of approximately 55 seconds. Several years ago, we showed that we could not replicate this observation. Recently, the original authors have claimed that we failed to find rhythms because 1) our flies did not sing enough and 2) our software for detecting song did not detect all song events. They claimed that they could detect rhythms in song annotated by hand. We report here that we cannot replicate their observation of rhythms in the hand-annotated data or in any dataset and that our original methods were not biased against detecting rhythms. We conclude that song rhythms cannot be detected.

## Introduction

When a male vinegar fly (*Drosophila melanogaster*) encounters a sexually receptive female, he performs a series of courtship behaviors, including the production of songs containing pulses and hums (or sines) via unilateral wing vibration (Fig. 1a). Every parameter of song displays extensive quantitative variation within a bout of singing, including the amplitude and frequency of pulses and sines and the timing of individual pulse and sine events (1, 2, 4–8). Like humans during conversation, *Drosophila* males modulate their song based on sensory feedback from their communication partner (4, 5).

**Figure 1.**
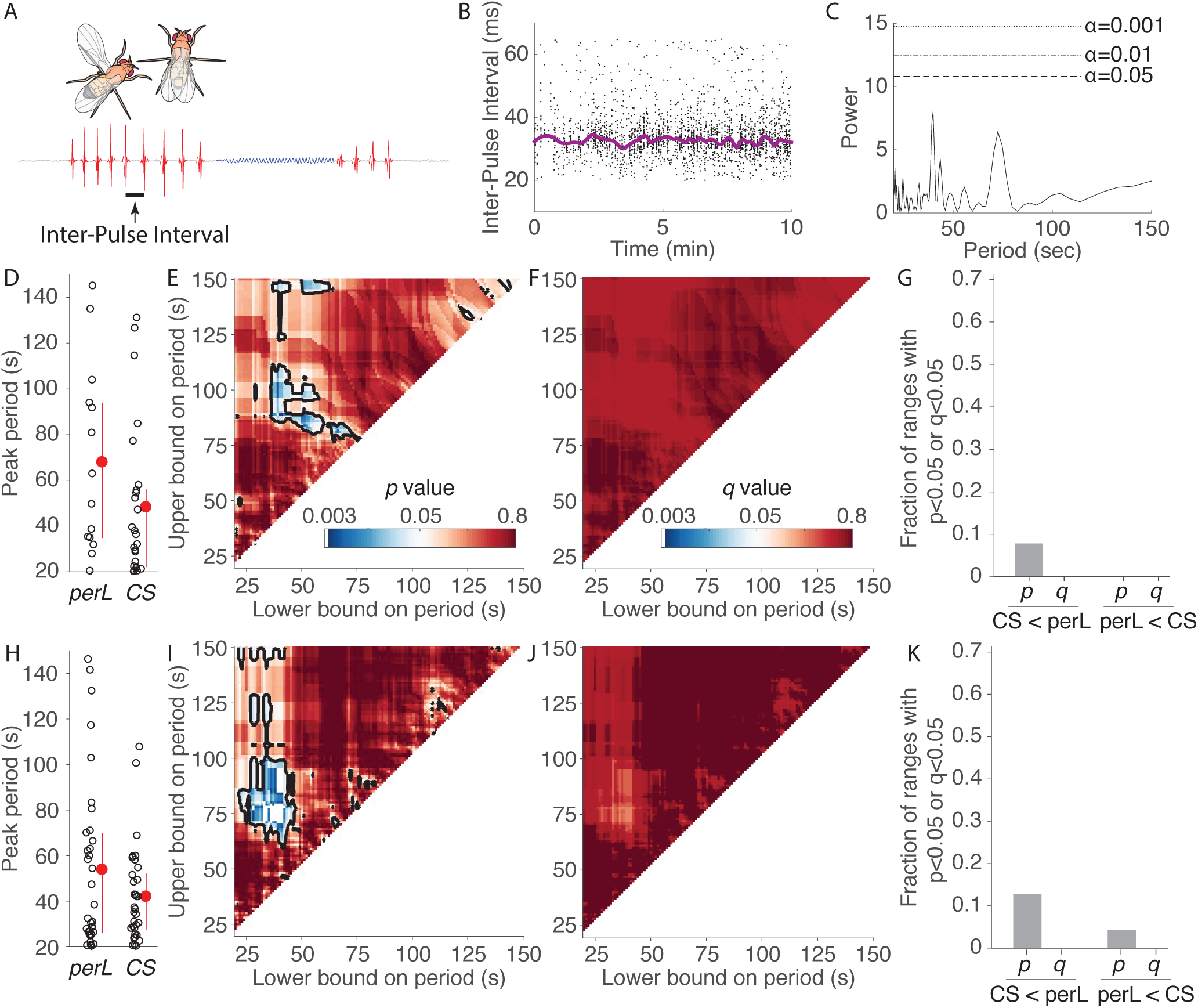
Genotype-specific periodicity cannot be detected in Drosophila courtship song. (A) Drosophila males produce courtship song, composed of pulses (red) and sines (blue), by extending and vibrating a wing. The inter-pulse interval is the time between consecutive pulses within a single train of pulses. (B) The average inter-pulse interval varies over time. (Purple line is rloess fit with sliding window of 200 samples). (C) Lomb-Scargle periodogram analysis of the inter-pulse interval data from panel (B) plotted for the range of 20 -150 sec. None of the peaks are significant at p < 0.05. (D) Comparison of the peak power between 20-150 sec from the Lomb-Scargle periodograms for the song data for the genotypes periodL (perL) and Canton-S (CS) manually-annotated by Kyriacou et al. (3). Red points and lines represent mean ± 1 SD for each genotype. (Right-tailed T-test p = 0.06. Rank Sum p = 0.10.) (E) P-values for period windows with different lower and upper bounds. (F) False discovery rate q values for the windows shown in (E). (G) Fraction of ranges with significant (p or q < 0.05) for either the test of Canton-S less than periodL or periodL less than Canton-S. (H-K) Same as (D-G) for newly collected song data from the same genotypes annotated using FlySongSegmenter. (H) (Right-tailed T-test p = 0.06. Rank Sum p = 0.45.)

Visual inspection of songs reveals that the mean inter-pulse interval varies over time (Fig. 1b). This observation was first made in 1980 by Kyriacou and Hall (10) and they reported that the mean cycled with a periodicity of about 55 sec and was controlled, in part, by the *period* gene, a gene required for circadian rhythms (11). Later papers demonstrated that evolution of a short amino-acid sequence within the *period* protein caused species-specific differences in this periodicity (11–14). These reports attracted considerable interest because they implicated the *period* gene in ultradian rhythms, in addition to its well-known role in circadian rhythms(15), and because it illustrated how genetic evolution can cause behavioral evolution.

Despite this progress, the molecular mechanisms causing this periodicity remained unknown. To further advance study of these rhythms, previously we searched for this periodicity using sensitive methods and failed to find evidence for song rhythms (1). We were mindful, however, that Kyriacou and Hall had argued that the presence or detectability of the rhythms was sensitive to assay conditions and methods of analysis (16). One of us, therefore, replicated the methods of Kyriacou and Hall as closely as possible, but, again, song rhythms could not be detected (2).

Kyriacou et al. (3) have recently questioned our previous conclusions. Here we focus on three major assertions that they claim call our conclusions into doubt. First, we examine their central claim that manual analysis of songs, but not automated analysis, reveals genotype-specific song rhythms. We find that re-analysis of their manually-annotated data provides no statistical support for genotype-specific rhythms. We also find no evidence for song rhythms in the original dataset and a new larger dataset. Second, we examined their claim that the original recordings contained insufficient data to detect rhythms and find that this claim is not supported by simulation studies. Third, we examine their claim that the high false negative rate of the automated song segmenter decreased the probability of detecting song rhythms and find no evidence that the missing pulse events biased our analysis of song rhythms. Further, we identify the major sources of false negative events in automated song analysis and illustrate that minor modifications to initialization parameters substantially improve performance of the song segmenter. Kyriacou et al. (3) also raised a number of minor concerns—such as how to choose an appropriate inter-pulse interval cutoff, whether temperature was controlled appropriately in our experiments, and whether songs produced beyond the first few minutes of courtship should be analyzed—that we consider peripheral to the central questions raised and therefore we have addressed these concerns (which are also unsupported by re-analysis) in the SI Appendix.

## Results

Earlier papers that identified song cycles employed several unusual methods of data analysis that it is useful to review. First, continuous inter-pulse interval data were binned into 10 sec intervals. We reported previously that binning the data, together with the analysis of relatively short songs, creates peaks in spectrogram analysis that fall within an artificially narrowed frequency range, corresponding approximately to the frequency range originally reported for the periodicity, and reduces the significance of periodiogram peaks (2 and see below). Despite the fact that this procedure squeezes periodogram results into a narrow frequency range, few songs contained peaks reaching a significance level of p < 0.05 (four of 149 songs, Fig. 3a of (2)), strongly suggesting that these peaks represent signals that cannot be distinguished from noise. All of the previously reported “statistically significant” comparisons of different genotypes are derived from analysis of mainly non-significant periodogram peaks. In this re-evaluation, we do not discuss binning, but instead focus on other methodological issues.

### No evidence that manual song segmentation reveals genotype-specific song rhythms

Kyriacou et al.’s (3) core finding is that different genotypes displayed different periodic rhythms of the inter-pulse interval. This is also the most important discovery reported in earlier papers on this subject (11–13, 17). Kyriacou et al. (3) manually annotated recordings made by Stern (2) from a wild-type strain, *Canton-S*, and a strain carrying a *period* gene mutation, *per*^*L*^, for flies they categorized as singing “vigorously.” We re-analyzed these data and the automatically segmented data (2). Flies homozygous for *per*^*L*^ display circadian rhythms that are longer than normal (15) and earlier papers have reported that *per*^*L*^ confers longer periods on the inter-pulse interval rhythm (10–13). Kyriacou et al. (3) report a difference in the mean song period between *Canton-S* and *per*^*L*^ with the manually annotated data, but not with the automatically segmented data, suggesting that song cycles exist and display genotype-specific frequencies and that the automatically segmented data is biased against detecting the song rhythm.

Kyriacou et al. (3) used several methods to measure periodicity in the original time series, which we discuss in more detail in the next paragraph. For approximately 85% of these songs, these methods do not yield statistically significant signals in the frequency range of 20-150 sec. Because most songs do not yield statistically significant peaks, Kyriacou et al. (3) identified the peak with maximum power in the range of 20-150 sec for each song and compared these values between genotypes. This is an unorthodox approach to data analysis. It is equivalent to sampling outliers from a distribution of random noise and then performing further statistics with these data. Nonetheless, Kyriacou et al (3) detected genotype-specific song rhythms using this method and so, below, we accept this premise and investigate whether there is statistical support for genotype-specific rhythms in the data. We start by examining whether there is evidence for rhythms in individual songs.

The general model proposed for these song rhythms is that the inter-pulse interval varies, on average, with a regular periodicity (10). Therefore, it should be possible to detect this rhythmicity with appropriate methods of periodogram analysis. We have previously employed Lomb-Scargle periodogram analysis (18–20) because this method does not require evenly spaced samples and Kyriacou et al. (3) also adopted this method. For example, the Lomb-Scargle periodogram of the time series in Fig. 1b is shown in Fig. 1c. In this case, despite the obvious variation in inter-pulse interval values observed in Fig. 1b, there is no significant periodicity between 20 and 150 sec. Kyriacou et al. (3) also employed Cosinor (21) and CLEAN (22) for periodogram analysis. CLEAN does not produce a significance value for periodogram peaks, so it is difficult to interpret. We find that Cosinor exhibits a high false positive rate (SI Appendix, Fig. S1), and should be avoided for this type of analysis.

Kyriacou et al. (3) state that wild-type *D. melanogaster* songs exhibit periodicity between 20 and 150 sec. Previously they reported that rhythms occurred with 50 – 60 sec periodicity (10). Increasing the width of the periodicity window from 50-60 sec to 20-150 sec increases the probability of detecting significant periods, but, even given this wide frequency range, we observed that only 4 of the 25 manually annotated *Canton-S* songs and 3 of the 25 automatically segmented songs contained periodogram peaks that reached a significance level of P < 0.05. (When we binned data in 10 sec bins, these values declined to 0 of 25 manually annotated and 1 of 25 automatically segmented songs.) These significant peaks are not localized to any particular narrow frequency range (SI Appendix, Fig. S1).

One reason to study non-significant peaks would be if periodicity is weak and not detected reliably by periodogram analysis. This seems unlikely, since simulated song rhythms can be detected with high confidence ((1, 2) and see below). Nonetheless, if periodogram analysis is underpowered, then we expect to observe that the major peak in most songs should display nearly-significant periodicity. In fact, we observe that 72% of p-values are greater than 0.2 (SI Appendix, Fig. S2). There is therefore no evidence that songs contain weak periodicity.

An alternative possible reason to include non-significant periodogram peaks in downstream analysis is that the signal to noise of the periodicity is extremely low. An analogue in neuroscience is that neural signals sometimes cannot be detected with high signal to noise and that only by averaging over many trials of a stimulus presentation can a neural response be detected robustly. We therefore examined the power distribution averaged over all the results for each genotype. These plots are essentially flat, suggesting that there is no signal hidden in the fluctuations of individual periodograms (SI Appendix, Fig. S3).

Given these observations, further analysis of these data seems unwarranted. However, Kyriacou et al. (3) compared the maximum periodogram peaks between 20-150 sec for the *Canton-S* and *per*^*L*^ recordings and found that the manually-annotated data showed a statistically-significant difference in the mean period, although the automatically segmented data did not (Fig. 3d of Kyriacou et al. (3)). This is the key result of their paper. We therefore attempted to replicate this observation. For the manually annotated data from each song we identified the peak in the periodogram of maximum power falling between a period of 20 and 150 sec. In contrast to their published results, we found that the average of the periods with maximum power (most of which were not significant) was not significantly different at P < 0.05 between the genotypes *Canton-S* and *per*^*L*^ (Fig. 1d). We have no explanation for this discrepancy between our statistical analysis and theirs.

Since there is no biological or quantitative justification for the particular frequency ranges examined in any study, we wondered whether the results were sensitive to the frequency range examined. We explored a wide range of possible frequency ranges and found that the test statistic was sensitive to the precise frequency range selected (Fig 1e). Most frequency windows do not generate a statistically significant difference between the genotypes (Fig. 1e,g) and false discovery rate correction for multiple testing (23, 24) yields no frequency ranges with significant results (Fig 1f,g).

Thus, there is no support for the specific results reported by Kyriacou et al. (3) and there is no statistical support for defining song inter-pulse interval cycle periods as occurring within any particular window. Most importantly, our analysis indicates that genotype-specific analysis of non-significant periodogram peaks has no justification. It is difficult to reconstruct precisely what steps in the analysis led previous reports to identify statistically significant genotype-specific differences, but it is possible that previous studies may have serendipitously selected frequency ranges that yielded significant results and/or did not properly control for multiple testing.

### New data provide no evidence for genotypic specific song periodicities

While we could not reproduce results reported by Kyriacou et al (3), we decided to take their observation at face value as a preliminary result and to test directly whether genotype specific song rhythms could be detected in a new, expanded data set. We recorded song from 33 *Canton-S* males and 34 *period*^*L*^ males. We identified the strongest periodogram peak in the frequency range of 20-150 s for each song and found no significant difference between these genotypes (Fig. 1h). We then compared test statistics across a wide set of frequency ranges, as described above. We identified some frequency ranges that yielded significant results in the predicted direction (Fig. 1i), with *period*^*L*^ rhythms slower than *Canton-S* rhythms, but for three reasons we believe these results are spurious. First, and most importantly, none of these ranges are significant after false discovery rate correction (Fig. 1j). Second, multiple frequency ranges support the *opposite* conclusion, that *Canton-S* rhythms are slower than *period*^*L*^ rhythms (Fig. 1k). Third, the frequency ranges yielding significant comparisons only partially overlap with the ranges found for the original dataset (c.f. Figs. 1e & 1i). In conclusion, there is not only no evidence that song rhythms exist, there is also no evidence that reported genotype specific differences in a song rhythm exist.

Putative song cycles cannot be identified in most automatically segmented song (2) and, as we showed above, in most manually annotated song. In addition, when statistically significant periodicity is detected, the frequencies of this periodicity do not cluster in a specific frequency range, but instead are spread randomly across the entire frequency range examined (SI Appendix, Fig. S5; Fig. 4 of Stern (2)). Finally, no songs are significant after correcting for multiple comparisons (Fig. 1). All together, these results imply that the few *statistically* significant periodicities that can be found do not carry *biological* significance.

### No evidence that low-intensity courtship provided insufficient data to detect song rhythms

While we found no statistical evidence for the existence of song rhythms or of genotype specific rhythms, we feel it is important to rebut several other statements made by Kyriacou et al. (3). They state that rhythms can be detected only in *songs* produced by “vigorously” singing males and write: “sporadic songs could not possibly provide any test for song cycles.” It is not clear if they mean that rhythms can be detected only in songs with many pulses or that only *flies* that sing songs with many pulses (“vigorous singers”) produce rhythms. Kyriacou et al. (3) manually annotated songs from flies that they categorized as vigorous and we showed above that significant periodicity can be found in only a minority of these songs and that these significant values are not localized to a particular frequency range (SI Appendix, Fig. S1d). Therefore, it is unlikely that only *flies* that sing songs with many pulses produce periodicity. We therefore performed simulations to determine whether rhythms can be detected only in *songs* with many pulses.

We previously investigated songs from 45-minute courtship recordings that contained at least 1000 inter-pulse interval measurements (2). Kyriacou et al. (3) argued that more than 180 inter-pulse interval measurements per minute (or approximately 5000 events in a 45-minute recording) should be identified to allow identification of song rhythms. To examine this claim, we performed a statistical power analysis using songs with variable numbers of inter-pulse interval measurements, where statistical power corresponds to the proportion of times periodicity is detected in songs where periodicity has been artificially imposed on song data (Fig. 2). We started with six 45-minute recordings of *Canton-S* from Stern (2) that contained more than 10,000 inter-pulse interval measurements. None of these six songs yielded statistically significant power in the frequency range between 50 and 60 sec (the range originally defined to contain rhythms (10)) and one song produced a marginally significant peak at 31.7 sec (P = 0.04), which falls between 20 and 150 sec (the range used by Kyriacou et al. (3)). Figure 2d and 2e illustrate the inter-pulse interval data and periodogram for one of these songs. Therefore, these songs do not contain strong periodicity in the predicted range and can serve as a template to examine the power of Lomb-Scargle periodogram analysis to detect simulated rhythms imposed on these data.

**Figure 2.**
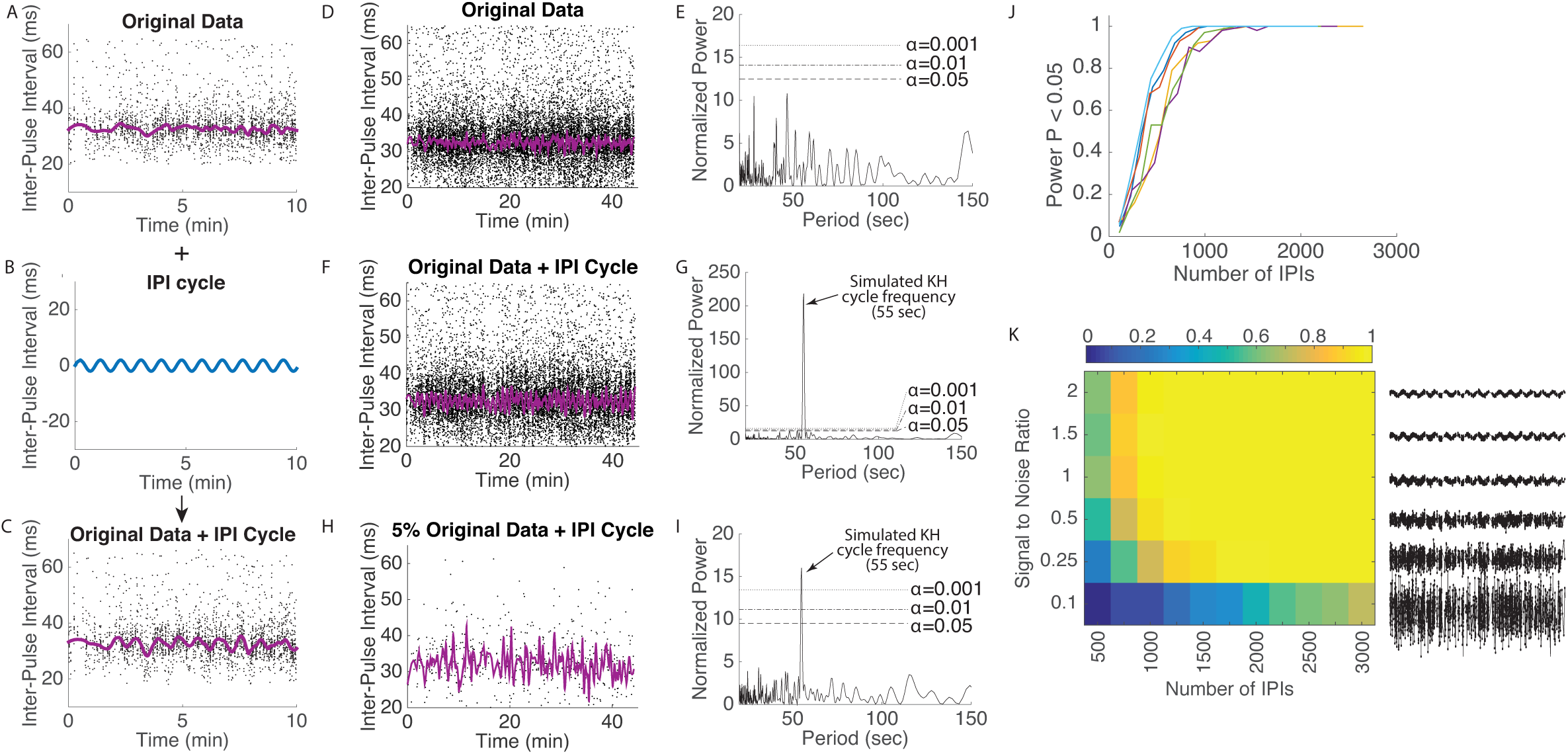
Simulations to explore power to detect rhythms, should they exist. (A-C) Example of how a periodic cycle was added to raw inter-pulse interval (IPI) data. Purple line in (A) illustrates the running mean of the raw data. Blue line in (B) shows a periodic rhythm with an amplitude of 2 msec and a period of 55 sec. Original data with simulated periodicity is shown in (C). (D) One example of 45 minutes of inter-pulse interval data. Purple line shows running mean. (E) Lomb-Scargle periodogram of data in (D) does not detect periodicity. (F) Data from (D) with a 55 sec periodicity imposed. (G) Lomb-Scargle periodogram of data in (F) now reveals a highly significant peak at 55 sec, consistent with the simulated periodicity. (H) Random removal of 95% of the inter-pulse interval data from (F). (I) Lomb-Scargle periodogram of the data in (H) detects significant periodicity. (J) Power analysis of six songs (each song a different color) containing more than 10,000 inter-pulse interval events after 55 sec periodicity was added and individual inter-pulse interval events were removed randomly. Power equals the fraction of times out of 100 that a song contained a rhythm with significant periodicity between 50 and 60 sec at P < 0.05. (K) Power to detect simulated noisy periodicity versus number of IPIs remaining after random removal of IPIs. Means of simulations for six songs containing more than 10,000 inter-pulse interval measurements are shown. Examples of simulated noisy rhythms are shown to the right. Colorbar shows power to detect simulated rhythm.

**Figure 3.**
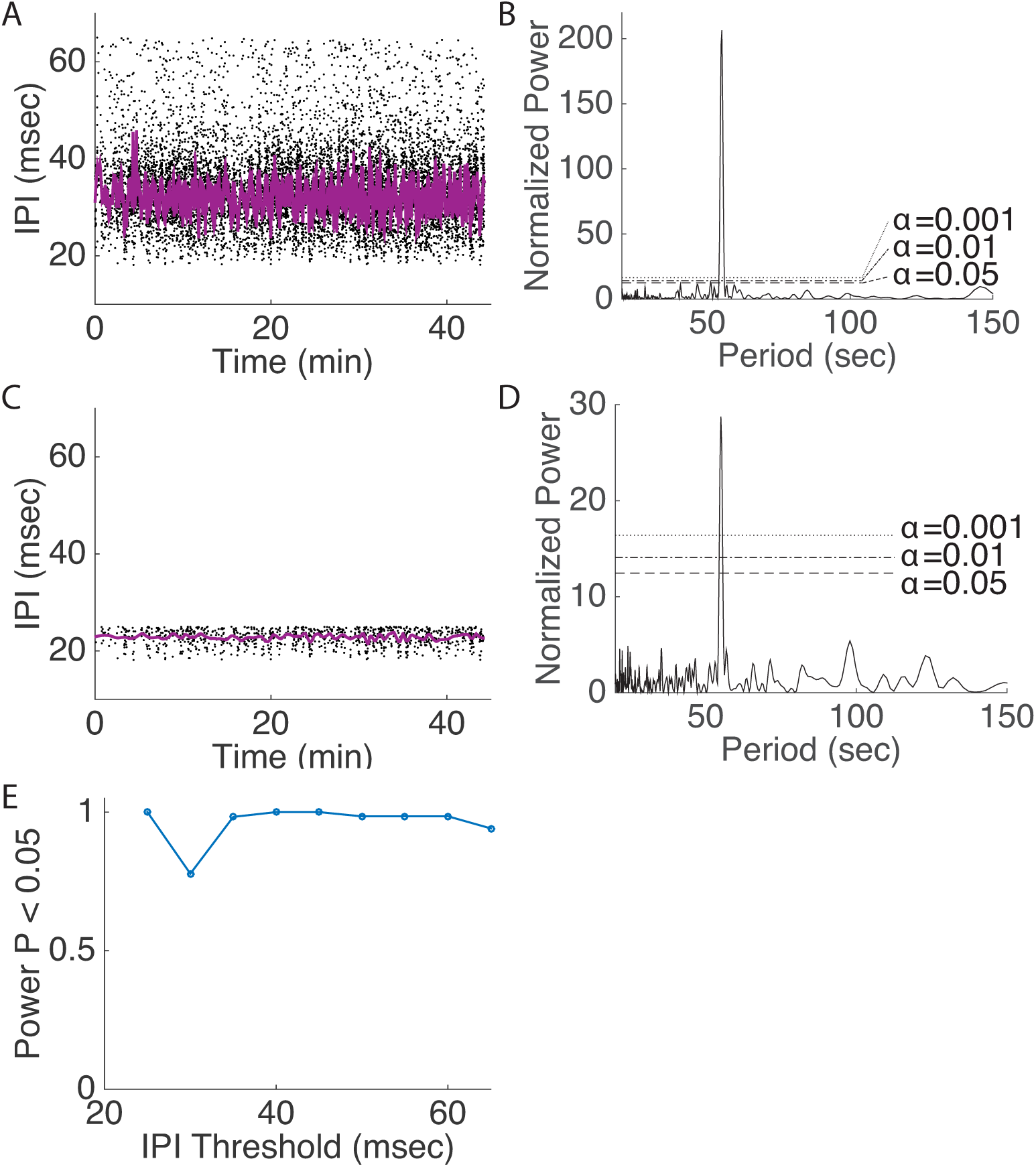
The specific inter-pulse interval threshold does not influence the statistical power to detect putative song rhythms. (A) Example of one original song with 55 sec periodicity artificially imposed on the original inter-pulse interval data. (B) Lomb-Scargle periodogram of data in panel (A), revealing strong signal at 55 sec. (C) Same simulated data as in panel (A) with all inter-pulse interval values greater than 25 sec removed. (D) Lomb-Scargle periodogram reveals strong signal of the simulated periodicity at 55 sec, even though the data were thresholded at 25 sec. (E) Power to detect simulated periodicity versus inter-pulse interval threshold for songs retaining at least 1000 inter-pulse interval values after thresholding.

The initial reports of periodic cycles in the inter-pulse interval reported rhythms with a mean period of 55 sec and an amplitude of approximately 2 ms (10). Therefore, we imposed a 55 second rhythm with an amplitude of 2ms on the six songs containing more than 10,000 inter-pulse interval measurements (Fig. 2a-c). We detected the simulated 55 sec rhythm in all six songs with P-values < 10e-74 (example shown in Fig. 2f,g). We then randomly removed data points from the songs iteratively and calculated the fraction of times we could detect the simulated rhythm with P < 0.05. We removed data randomly from the dataset to simulate the effect of failing to detect individual events in the song and we also removed chunks of data (in 10 sec bins) to simulate large gaps between song bursts, such as might be generated during low-intensity courtship. We found that in both scenarios we could randomly remove at least 90% of the data and still detect simulated rhythms at least 80% of the time (example shown in Fig. 2h,i; summary statistics shown in Fig. 2j and SI Appendix, Fig. S4a). That is, as long as songs contained at least 1000 inter-pulse interval measurements, Lomb-Scargle periodogram analysis detected simulated rhythms with power greater than 0.8. Similar results were found when we analyzed only the first 400 sec of songs (SI Appendix, Fig. S4c,d). Furthermore, periodicity could be detected with power greater than 0.8 when the amplitude of simulated periodicity was greater than at least 1 msec (SI Appendix, Fig. 4b). These results were robust to noise in the original periodicity. Song with a signal to noise ratio of as low as 0.25 could be detected with power > 0.7 with sample sizes of at least 1000 inter-pulse interval measurements (Fig. 2k). Similarly, periodicity could be detected reliably when we simulated a non-sinusoidal rhythm (SI Appendix, Fig. Fig. S4e) and when periodicity was imposed for only a fraction of the total song (SI Appendix). Thus, Lomb-Scargle periodogram analysis is a sensitive method for detecting simulated periodicity, even in the presence of noise or discontinuities in the waveform.

Songs containing at least 1000 inter-pulse intervals provide sufficient data to identify putative song cycles. In fact, we find that songs can be deeply corrupted by the absence of large segments of song and simulated periodicity can still be detected.

### No evidence that the automated fly song segmenter biased the results

Kyriacou et al. (3) expressed concern that our automated fly song segmenter displayed a low true positive rate (the segmenter failed to detect approximately 50% of the pulses identified through manual annotation) and produced some false positive calls (approximately 4% of events scored as pulses by the automated segmenter appear to be noise). They suggest that these incorrect pulse event assignments could bias estimation of the mean inter-pulse interval and therefore decrease the signal-to-noise of the periodic cycle, making it difficult to detect a periodic signal. In principle, a large sample of incorrect calls could bias results, so we investigated whether this was the case for our prior analyses. We used Kyriacou et al.’s (3) manually-annotated dataset to investigate the potential for bias and to evaluate performance of the automated segmenter.

When a single pulse event is not detected, the inter-pulse interval is then calculated as the sum of the two neighboring real intervals. On average, this is approximately double the average inter-pulse interval. The average inter-pulse interval for the *Canton-S* recordings reported in Stern (2) is approximately 35 msec with a standard deviation of approximately 7 msec. Therefore, skipping a single pulse event is expected to result in inter-pulse interval measurements of approximately 70 msec, but with considerable variance. Following Kyriacou and Hall (16), Stern (2) employed a heuristic threshold of 65 msec to reduce the number of spurious inter-pulse interval values. Therefore, in the specific case when a single pulse in a train is missed, approximately one third of the incorrectly scored doublet inter-pulse interval measurements would be shorter than 65 msec and are expected to contaminate the original dataset.

However, this scenario applies only when one undetected pulse is flanked by two pulses that are detected. Skipping more than one pulse would always result in inter-pulse interval measurements that are excluded by the 65 ms threshold. We found, however, that only 9% of the pulses missed by automated segmentation were singletons (SI Appendix, Fig. S6a). These incorrect inter-pulse intervals contribute to a slight excess of inter-pulse intervals with high values (SI Appendix, Fig. S6b). Lowering the inter-pulse interval threshold would, therefore, remove most or all spurious inter-pulse intervals. Since our power analysis, discussed above, revealed that periodogram analysis was robust to random removal of inter-pulse interval events, as long as songs still contained at least 1000 values, loss of a small number of inter-pulse intervals is not expected to hamper detection of rhythms. After reducing the inter-pulse interval threshold to 55 msec, we still found no compelling evidence for significant periodicity in the original data (SI Appendix, Fig. S7). Therefore, we explored the effect of reducing the inter-pulse interval cutoff even further. In this case, we used all 68 *Canton-S* songs from Stern (2) and retained for analysis only those songs that contained at least 1000 inter-pulse interval measurements after imposing the new inter-pulse interval threshold. We explored a range of cutoff values from 25 to 65 msec. We found that we could detect the simulated rhythm in most songs with at least 1000 inter-pulse interval measurements remaining after thresholding, even when the threshold was as low as 25 msec (Fig. 3). Therefore, we can find no evidence that pulses missed by the automated song segmenter or the specific inter-pulse interval threshold used in Stern (2) prevented detection of song rhythms.

Although detection of putative song rhythms is robust to dropped pulses in songs that retain at least approximately 1000 inter-pulse intervals, it is worth reviewing briefly why the segmenter failed to detect certain pulses in recordings reported in Stern (2). The first step of song segmentation involves detection of pulse-like signals and sine-like signals (1). In subsequent steps, the segmenter filters out many kinds of sounds that were originally classified as song pulses. Both the initial detection of pulses and subsequent filtering steps are sensitive to multiple parameters. These parameters are specified prior to segmentation and can be modified to enhance performance of the segmenter for different recordings. We identified two primary causes for missed pulses. First, Stern (2) recorded song in larger chambers than those used previously with these microphones (1), to match the chamber size used by Kyriacou & Hall (10). This larger chamber with one microphone had reduced sensitivity compared to the original smaller chamber. The segmenter thus tended to miss pulses of lower amplitude, which are hard to automatically differentiate from noise, and this explains approximately 35% of the missed pulses (SI Appendix, Fig. S8a, c).

The second major cause of missed pulses is that *Drosophila* males produce pulses with a range of carrier frequencies (tones). The higher frequency pulses tend to resemble other non-song noises, like grooming, and a user can set parameters in the segmenter to attempt to exclude these non-song noises based on the carrier frequency of the event. Stern (2) used parameters to minimize the false positive rate, including a relatively low carrier frequency cutoff for pulses. The lower pulse frequency threshold used by Stern (2) explains approximately 42% of the missed pulses (SI Appendix, Fig. S8b,d). Using the same software with different parameters (from Coen et al. (5)) recovers many of these high-frequency pulses without substantially increasing the false positive rate (SI Appendix, Fig. S8c-f).

Above, we showed that including more pulse events, by manual annotation, did not increase the probability of detecting song rhythms. Therefore, there is no evidence that the data resulting from the song segmenter parameters used in Stern (2) generated a data set that was biased against detection of song rhythms. While the song segmenter does not detect all pulse events that can be detected by manual annotation, the segmenter does provide data sets that are several orders of magnitude larger than those that can be generated by manual annotation, which has allowed discovery of multiple new phenomena related to *Drosophila* courtship song (4–6). In addition, the sensitivity of the song segmenter can be improved with optimization of initial parameters, as expected of any segmentation algorithm.

## Discussion

We cannot detect a periodic cycling of the inter-pulse interval in *Drosophila* courtship song even in the songs manually annotated by Kyriacou et al. (3) and used as evidence for periodicity in their paper. While it is impossible to prove a negative, our results agree with previous analyses that have concluded that there is no statistical evidence that these rhythms exist (1, 2). In particular, by exploring some of the relevant parameter space with statistical tests on the song that was manually-annotated by Kyriacou et al. (3), we find that subsets of parameters sometimes produce p-values lower than 0.05, but that (1) few regions of parameter space generate “significant” results, (2) these “significant” regions are scattered apparently randomly in parameter space, and (3) none of these “significant” results survive multiple test correction (Fig. 1).

Previously, we offered one explanation for how apparent song rhythms may have been detected. We found that binning data from short songs confined the periodogram peaks with maximum power close to the range reported as the song cycle (2). While few of these peaks reached statistical significance, previous authors have accepted these peaks as “signal” and performed statistical analyses to compare the peaks between genotypes. All “statistically significant” results from earlier papers were derived mainly from non-significant peaks in periodogram analysis and from relatively small sample sizes (usually fewer than 10 flies of each genotype), so it is questionable whether these derivative statistics are valid. Genotype-specific periodicities reported in earlier papers may have resulted, by chance, from studies of a small number of short songs that fortuitously led to occasional apparent replication of the original observations.

There may be a more prosaic explanation for the initial discovery of song cycles. Every fly produces highly variable inter-pulse intervals. In addition, a running average of these data reveals that the average inter-pulse interval cycles up and down (Fig. 1b), similar to the temporally-binned data first reported by Kyriacou and Hall (10). There is no debate about this observation. The claim in dispute is that the average inter-pulse interval cycles regularly. We can find no evidence for this claim. It is easy to imagine, however, that visual examination of short recordings of song would make it appear as if the mean inter-pulse interval cycled regularly.

The extraordinary within-fly variation in the inter-pulse interval and in the mean inter-pulse interval may result from multiple causes, including the possibility that male flies respond to ever-changing cues during courtship and modulate their inter-pulse interval to optimize their chances of mating. Individual *Drosophila* males modulate specific aspects of their courtship song based on their own patterns of locomotion and in response to feedback from females, including the transition between sine and pulse song (5) and the amplitude of pulse song (4). There is additional evidence that males modulate the carrier frequency of sine song (1). We hypothesize that male flies also modulate their inter-pulse interval in response to specific internal or external cues.

We can find no statistical evidence for periodicity of the inter-pulse interval in individual courtship songs and no evidence that comparisons of the strongest periodogram peaks from each song identify genotype-specific rhythms. These results hold *both* for the songs manually annotated by Kyriacou et al. (3) and for two independent large datasets automatically annotated with FlySongSegmenter using optimized parameters. At this time, a conservative assessment of the problem is that *Drosophila* courtship song rhythms and genotype-specific effects on these rhythms cannot be replicated.

## Methods

Courting fruit flies of Oregon-R and *per*^*L*^ were recorded as described previously (2). All analyses were performed in Matlab. All data and code are freely available, as described in the Software and Data Availability section. Further methods can be found in SI Appendix.

## Acknowledgements

We thank Elizabeth Kim for recording the new samples of flies.

### Software and data availability

Computer code for all analyses described in this paper is available at https://github.com/murthylab/noIPIcycles. Code for the version of FlySongSegmenter used in Cohen et al. (5) is available at https://github.com/murthylab/songSegmenter. The raw and segmented song data for the new song recordings is available at https://www.janelia.org/lab/stern-lab/tools-reagents-data.

